# Investigation of swine caecal microbiomes in the northern region of Thailand

**DOI:** 10.1101/2023.07.03.547425

**Authors:** Thanaporn Eiamsam-ang, Pakpoom Tadee, Songphon Buddhasiri, Phongsakorn Chuammitri, Nattinee Kittiwan, Ben Pascoe, Prapas Patchanee

**Affiliations:** Graduate Program in Veterinary Science, Faculty of Veterinary Medicine, Chiang Mai University, Muang, Chiang Mai, Thailand; Integrative Research Center for Veterinary Preventive Medicine, Faculty of Veterinary Medicine, Chiang Mai University, Muang, Chiang Mai, Thailand; Department of Veterinary Biosciences and Public Health, Faculty of Veterinary Medicine, Chiang Mai University, Muang, Chiang Mai, Thailand; Veterinary Research and Development Center (Upper Northern Region), Hang Chat, Lampang, Thailand; Big Data Institute, University of Oxford, Oxford, United Kingdom; Department of Biology, University of Oxford, Oxford, United Kingdom; School of Animal and Comparative Biomedical Sciences, University of Arizona, Tucson, AZ, USA

**Keywords:** Swine, commercial swine farming, shotgun metagenomics, caecal microbiome, antimicrobial resistance (AMR)

## Abstract

**Introduction:** The northern region of Thailand serves as a crucial area for swine production, contributing to the global food supply. Previous studies have highlighted the presence of foodborne pathogens originating from swine farms in this region, posing a threat to both human and animal health.

**Gap statement:** Multiple swine pathogens have been studied at a species level, but the distribution and co-occurrence of pathogens in agricultural swine has not been well established.

**Aim:** Our study employed the intestinal scraping technique to directly examine the microorganisms interacting with the swine host.

**Methodology:** We used shotgun metagenomic sequencing to analyse the caecal microbiomes of swine from five commercial farms in northern Thailand.

**Results:** Swine caecal microbiomes contained commensal bacteria such as *Bifidobacterium*, *Lactobacillus*, and *Faecalibacterium*, which are associated with healthy physiology and feed utilisation. We also identified multiple pathogenic and opportunistic bacteria present in all samples, including *Escherichia coli*, *Clostridium botulinum*, *Staphylococcus aureus*, and the *Corynebacterium* genus. From a One Health perspective, these species are important foodborne and opportunistic pathogens in both humans and agricultural animals. Antimicrobial resistance genes were also detected in all samples, specifically conferring resistance to tetracycline and aminoglycosides which have historically been used extensively in swine farming.

**Conclusion:** The findings further support the need for improved sanitation standards in swine farms, and additional monitoring of agricultural animals and farm workers to reduce contamination and improved produce safety for human consumption.

## Introduction

The northern region of Thailand serves as a crucial area for swine production, contributing to the global food supply. In 2022, swine production, export, and domestic consumption in Thailand were 1.45 million tons, 5,676 tons, and 1.15 million tons, respectively. The pork industry in Thailand is vital for protein and meat production, with quality control standards beginning at the farm level (The Swine Raisers Association of Thailand, 2023). Furthermore, northern swine farms have diverse farming types and farm scales (Department of livestock and development, 2023). Previous studies have highlighted the presence of foodborne pathogens originating from swine farms in this region, posing a threat to both human and animal health (Patchanee et al., 2020). Gut microbiomes include the full complement of microorganisms that live in the digestive tracts of living animals (Xiao et al., 2016) and diversity in the gut microbiome reflects variation in the host species, which can be influenced by farming conditions, environment, and antimicrobial usage (Looft et al., 2014; Guevarra et al., 2018; Quan et al., 2020).

Culture-independent shotgun metagenome sequencing can yield data on the co-occurrence of bacterial species (Hugenholtz et al., 1998; Quan et al., 2020). By comparing information on the microbial diversity of the gut microbiome, antimicrobial resistance genes, and genes involved in other biological and metabolic processes, we can better investigate the relationship between microbes and swine (Looft et al., 2014; Chen et al., 2017). Furthermore, this technique can be used to trace pathogen transmission from the swine gastrointestinal tract to humans, encompassing One Health principles (Kim et al., 2021; Duarte et al., 2021; World Organization for Animal Health, 2023). Investigations of differences in the microbiomes of livestock animals have included studies on the impact of antimicrobial growth promoters on microbial abundance and metabolic profiles in poultry (Zou et al., 2022), and altered nutrient utilisation and secondary metabolic production in cows (Strachan et al., 2023).

In swine, shotgun metagenomic analyses have also been applied to investigate the antimicrobial and heavy metal resistance genes harboured in fattening swine caecal microbiomes, as well as the effect probiotic supplementation has on piglet health (Tunsagool et al., 2021; Apiwatsiri et al., 2022). While previous studies have shown that caecal microbiomes are diverse and can be correlated with high feed efficiency and utilisation (Quan et al., 2020), few studies have used the intestinal scraping technique to study microorganisms directly interacting with the host, particularly in swine. In this study, we employed the intestinal scraping technique to directly examine the microorganisms interacting with the swine host from commercial farms in the northern region of Thailand. Shotgun metagenomic sequencing was used to characterise the swine gut microbiome, focusing on bacterial diversity, genes related to the metabolism, and antimicrobial resistance genes.

## Materials & Methods

### Swine caecal sampling

Five swine caecal samples from five commercial swine farms in northern Thailand were included in this study. All samples were collected from farms located in the northern region of Thailand, including Lampang, Chiang Mai, Chiang Rai, and Uttaradit provinces. All farms represented medium- (between 500–5000 pigs per farm) and large-scale (more than 5,000 pigs per farm) commercial farms (**Figure 1**). The market-weighted swine from five commercial farms (4 medium and 1 large farms size) were slaughtered at 120 days of age. The caeca were collected during the period March 2021 through July 2021 at the slaughtering process and stored at -30 ^°^C. Subsequently, the caecal contents were collected using the intestinal scraping techniques with the purpose of studying microorganisms that directly interact with the host. Experimental protocols were approved by the Animal Care and Use Committee (FVM-ACUC), Faculty of Veterinary Medicine, Chiang Mai University (Ref. No. S2/2564).

**Figure 1:**
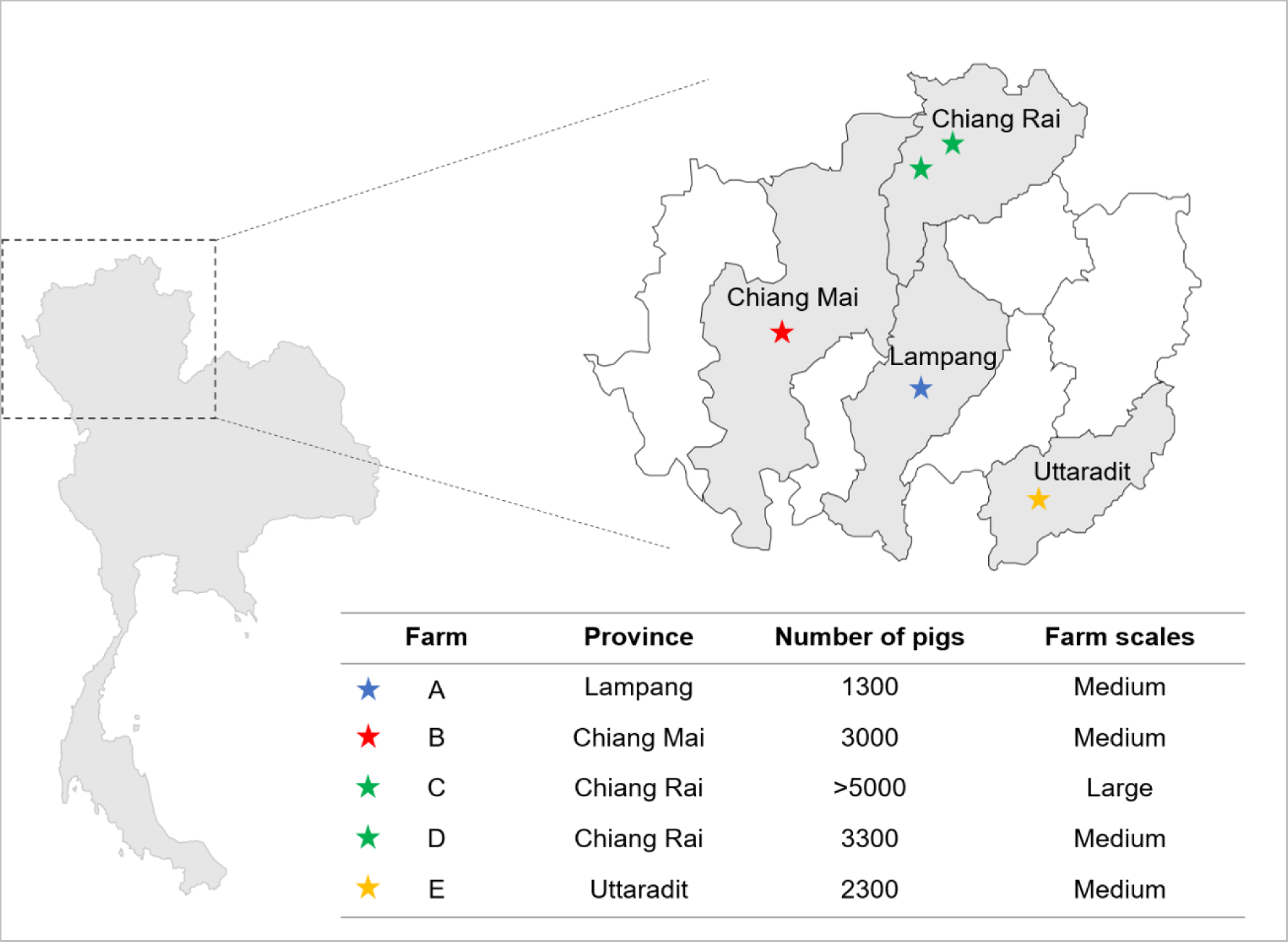
Geographical distribution and the details of five commercial swine farms in this study.

### DNA extraction and shotgun metagenomic sequencing

Genomic DNA was extracted from caecal tissues using a Tissue Genomic DNA Extraction Kit (Qiagen). UV-Vis spectrophotometry and 1% agarose gel electrophoresis were used to determine DNA concentration and quality. Paired-end library preparation with read length 150 bp was performed using the TruSeq Nano DNA Kit (Illumina), and libraries were sequenced on the Illumina NovaSeq 6000 platform. The total reads obtained from the metagenomic sequencing was 138,868,453 reads (an average of 27,773,691 reads per sample) (**Supplementary Table S1**). Data can be accessed at the National Center for Biotechnology Information with accession number PRJNA937456 (individual accession numbers in **Supplementary Table S2**).

### Data quality control and assembly

Sequencing reads were first screened by FastQC (version 0.11.9) for quality assessment (Andrew et al., 2010) and CutAdapt used to trim low-certainty bases and poor-quality reads (Martin, 2011). Reads were mapped to the swine reference genome (Sscrofa11.1; BioSample ID SAMN02953785) to remove host genome contamination using bowtie2 (v2.3.3) and Samtools (v.1.12) (Langmead & Salzberg, 2012; Danecek et al., 2021). Short reads were assembled *de novo* into contigs with metaSPAdes using k-mer sizes of 21, 33, 55 and 77 (Nurk et al., 2017). Contigs were binned using an expectation-maximization algorithm in MetaBAT2 with a minimum contig length of 1500 (Kang et al.,2019). MetaGeneMark was used to predict potential ORFs (Zhu et al., 2010). Gene sequences with less than 100 bp in length were removed, and remaining sequences were translated into the corresponding amino acid sequences. All predicted genes were clustered, and redundancy was eliminated using CD-HIT (identity > 95%, coverage > 90%) (Fu et al., 2012).

### Taxonomic classification of the swine caecal microbiome

The taxonomic classification of the assembled contigs was performed using Kraken2 against a standard database with the default parameters (Wood et al., 2019). The estimate of relative abundance from Kraken2 was determined using Bracken (Lu et al., 2017). The taxonomic abundance of bacterial composition in each sample was visualized as the balloon plot and the 100% stacked bar graph by ggpubr and ggplot2 packages in R (Team R, 2020).

### Functional annotation and profiling of the swine caecal microbiome

For the functional annotation of the swine caecal microbiome, the putative amino acid sequences of five swine caecal metagenomic contigs acquired from MetaGeneMark were functionally annotated and profiled according to Kyoto Encyclopedia of Genes and Genomes (KEGG) databases using GhostKOALA (Kanehisa et al., 2016). Obtained KEGG Orthology were reconstructed to the KEGG pathway and classified into metabolic pathway at level 1, 2 and 3. The visualization of the KEGG categories and the KEGG pathway in each level were performed using the balloon plot and the 100% stacked bar graph by ggpubr and ggplot2 packages in R (Team R, 2020).

### Identification of antimicrobial resistance genes

Assembled contigs were used to detect antimicrobial resistance genes harboured by isolates in the swine caecal microbiome using ABRicate against the CARD database (CARD: The Comprehensive Antibiotic Resistance Database) with DNA identity > 85% and coverage > 80% (Seemann T., 2016; Alcock et al., 2020). The detected antimicrobial resistance genes (ARGs) were visualised as a binary heatmap and a Venn diagram which constructed using heatmaply and ggplot2 packages in R (Team R, 2020).

## Results

### Bacterial taxonomic diversity in swine caecal microbiomes

Large- (more than 5,000 pigs per farm) and medium-scaled (between 500-5,000 pigs per farm) swine farms were selected from the northern region of Thailand to assess the composition of swine caecal microbiomes (**Figure 1**). Caecal samples from different provinces were extracted and sequenced using a shotgun metagenomics approach on an Illumina Novaseq 6000 platform, which generated an average of 27,112,029 quality filtered reads per sample. After host genomic removal, the remaining reads were an average of 9,221,845 reads per sample. Each individual sample’s sequence reads were assembled *de novo*. The number of contigs ranged from 65,294 to 229,191 contigs, and the assembled contigs were used to classify the bacterial taxonomic composition of each swine caecal sample (**Supplementary Table S1**).

Based on the relative abundance, the bacterial phylum-level composition was diverse among the samples. The *Firmicutes* were the most prevalent phyla in swine caeca from farms B, C, and E; while *Actinobacteria* constituted the most common phylum in swine caeca from farm A and *Proteobacteria* was the most common phylum in swine caeca from farm D. Farms C and D were located in the same province (Chiang Rai province), but the relative phylum-level composition in swine caeca was different, with farm C containing a higher abundance of *Firmicutes* and *Bacteroidota* phyla than farm D. In contrast, swine caeca from farm D were comprised with a higher proportion of *Proteobacteria* Phyla than farm C. Bacterial phyla identified in swine caeca from farm E were the most distinct compared with other farms. Farm E was comprised of a higher proportion of *Bacteroidota* and a lower proportion of *Actinobacteria* than other farms (**Figure 2**).

**Figure 2:**
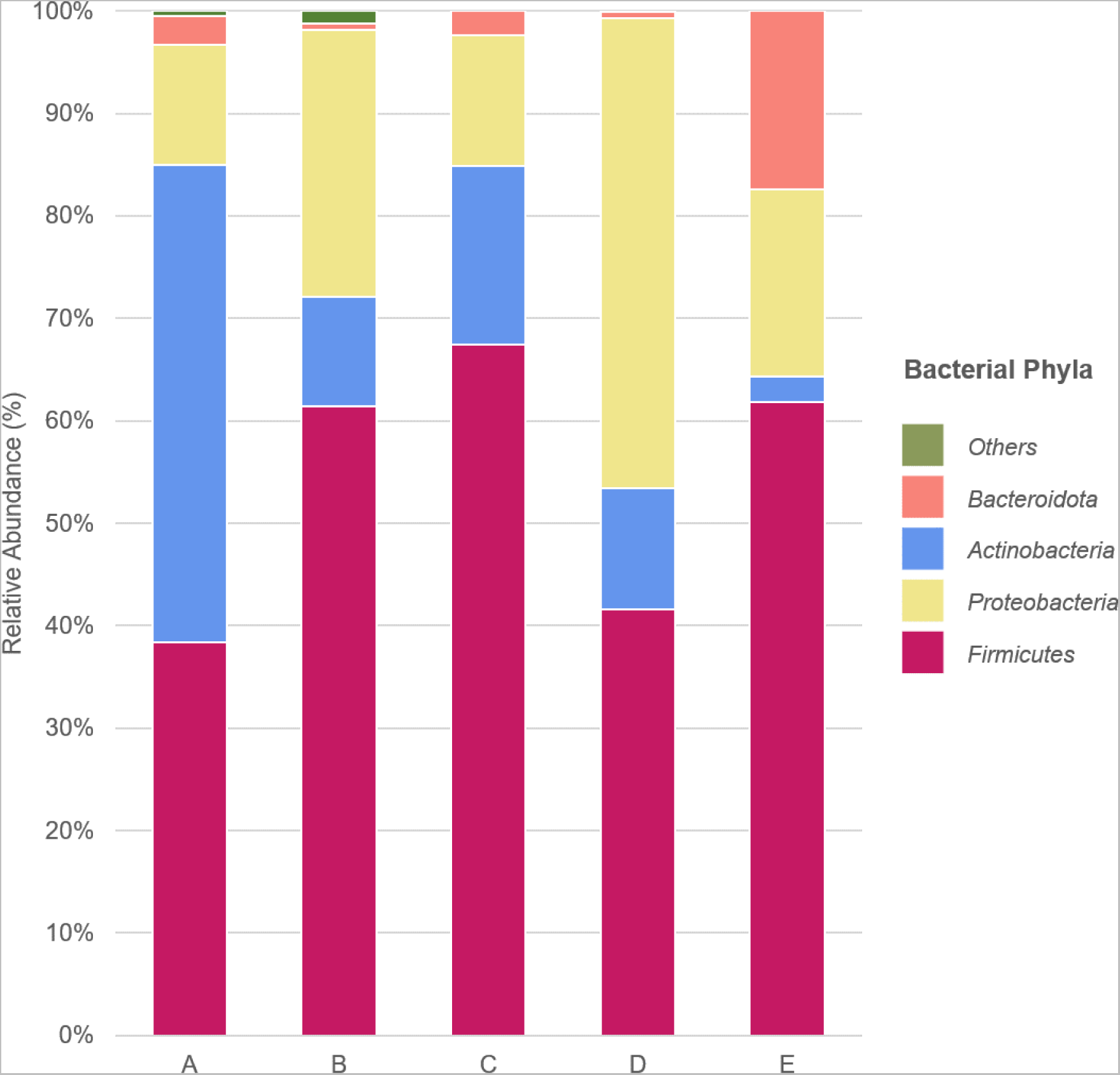
The relative abundance and diversity of the swine caecal microbiome at phylum level distributed in each commercial farm.

Considerable diversity was also observed in the relative proportion of bacterial genera identified in swine caeca from each farm. *Escherichia*, *Clostridium*, and *Faecalibacterium* were the most common genera identified in caeca from all farms. Different bacterial genera were dominant in caeca from different farms; *Corynebacterium* was the dominant genera found in caeca from farm A, followed by *Escherichia*. *Escherichia* was the most common genus found in caeca from farm B, followed by *Clostridium, Staphylococcus* and *Streptomyces*. *Lactobacillus* and *Bifidobacterium* were the most common genera in caeca from farm C and *Collinsella* was the predominant genera found in caeca from farm D. Both *Lactobacillus* and *Escherichia* genera were found in the same proportion in caeca from farm E, where the *Bacteroides* and *Prevotella* genera (both *Bacteroidota* phyla) were also common (**Figure 3** and **Supplementary Figure S1**).

**Figure 3:**
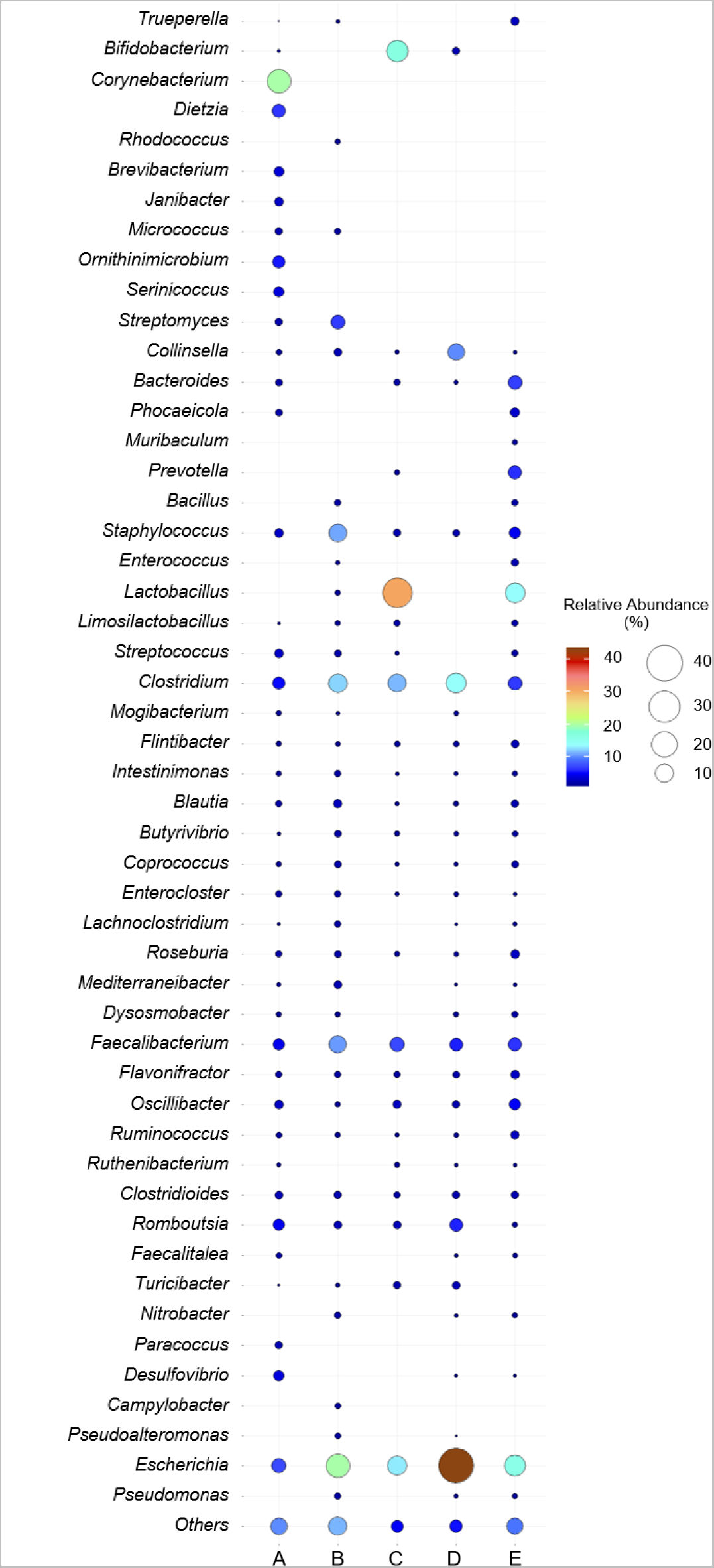
The bacterial taxonomic balloon plot of the relative abundance and diversity of the swine caecal microbiome at genus level distributed in each commercial farm. Circle sizes and color represented the percentage of the relative abundance.

At the species level, swine caeca from each farm contained a diverse range of pathogenic, opportunistic, and commensal bacteria. *Escherichia coli*, *Clostridium botulinum*, and *Faecalibacterium prausnitzii* were the predominant species in all samples. Similar to the genera-level composition, there are some predominant bacterial species found in each individual farms including, *Corynebacterium urealyticum* and *Corynebacterium xerosis* in caeca from farm A while *Streptomyces hygroscopicus* and *Streptomyces sp. HF10* were the predominant species and found only in caeca from farm B. Correlated with the genera level, probiotic species *Lactobacillus amylovorus, Bifidobacterium choerinum, Bifidobacterium pseudolongum,* and *Bifidobacterium thermophilum* were the predominant species in caeca from farm C. Different species from the genus *Lactobacillus* were most common in caeca from farm E and C microbiomes; the predominant species in caeca from farm E was *Lactobacillus crispatus* and *Lactobacillus johnsonii*, which differed to the *Lactobacillus* species found in farm C. In Farm D, *Collinsella aerofaciens* was the predominant species, followed by *Escherichia coli* (**Figure 4** and **Supplementary Figure S2**).

**Figure 4:**
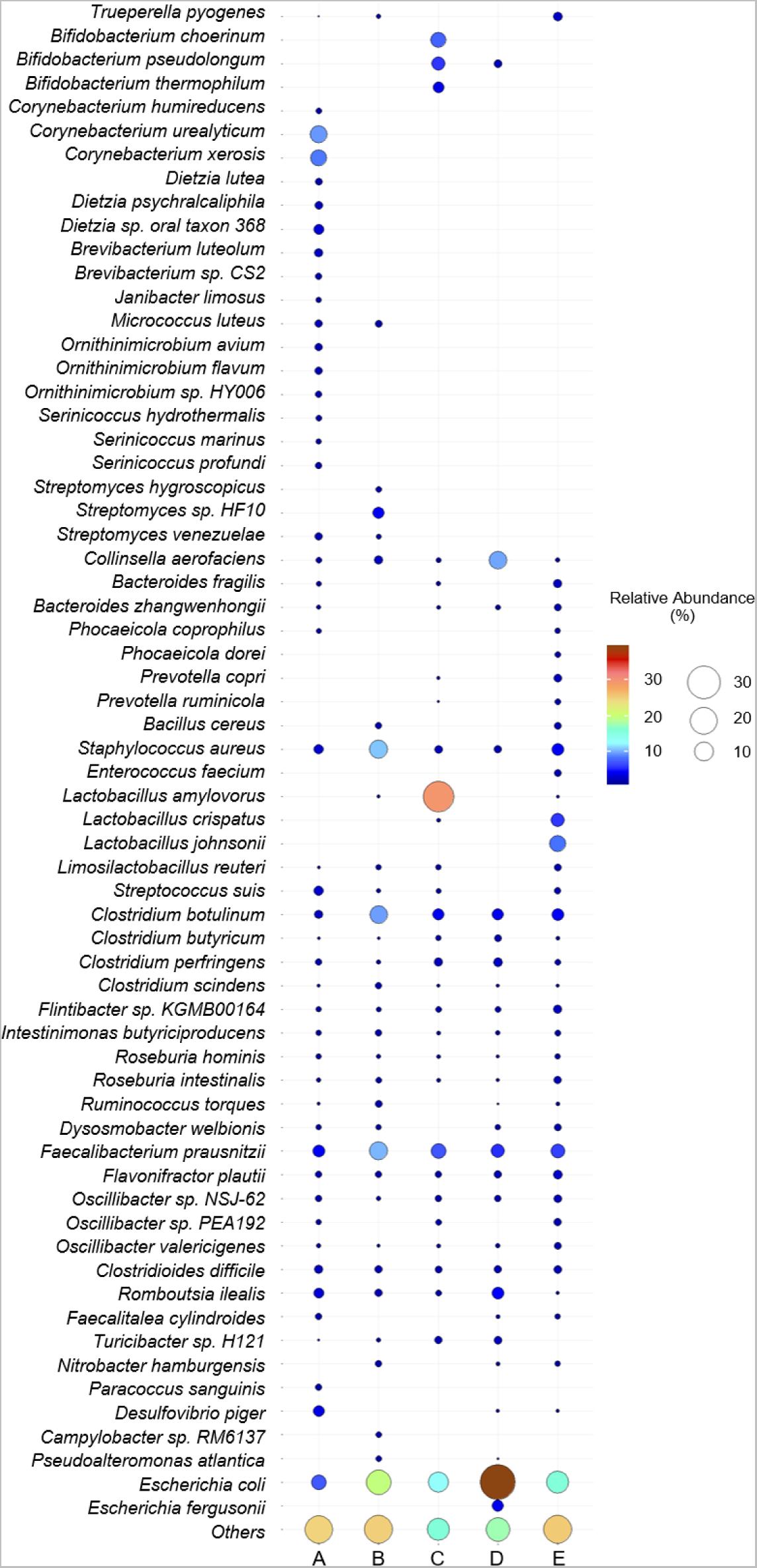
The bacterial taxonomic balloon plot of the relative abundance and diversity of the swine caecal microbiome at species level distributed in each commercial farm. Circle sizes and color represented the percentage of the relative abundance.

### Similarity in functional characterization of swine caecal microbiome genes identified between farms

The predicted genes were functionally annotated using the Kyoto Encyclopedia of Genes and Genomes (KEGG) databases to assess the potential functional capacity of the swine caecal microbiomes. In total, 3.8-12.3% of the predicted genes were assigned to the KEGG Orthology and reconstructed to the KEGG pathway (**Supplementary Table S1**). All swine caecal microbiome samples contained genes from all four main pathway categories. Genes from the KEGG pathway categories correlated with metabolism were the most commonly identified genes on all farms, followed by genetic information, environmental information, and cellular processes categories. There is quite a similar proportion of the relative abundance of the KEGG pathway genes identified between farms (**Figure 5**). According to the KEGG pathway at level 2, carbohydrate metabolism was the most prevalent metabolic pathway, followed by amino acid, energy, and nucleotide metabolism. Genes identified in microbiome samples from all farms included those with roles in replication, repair, and translation associated with the genetic information process. Furthermore, gene-related with environmental information and cellular processes including membrane transport, signal transduction and cellular community-prokaryotes were also found among all samples (**Figure 5**).

**Figure 5:**
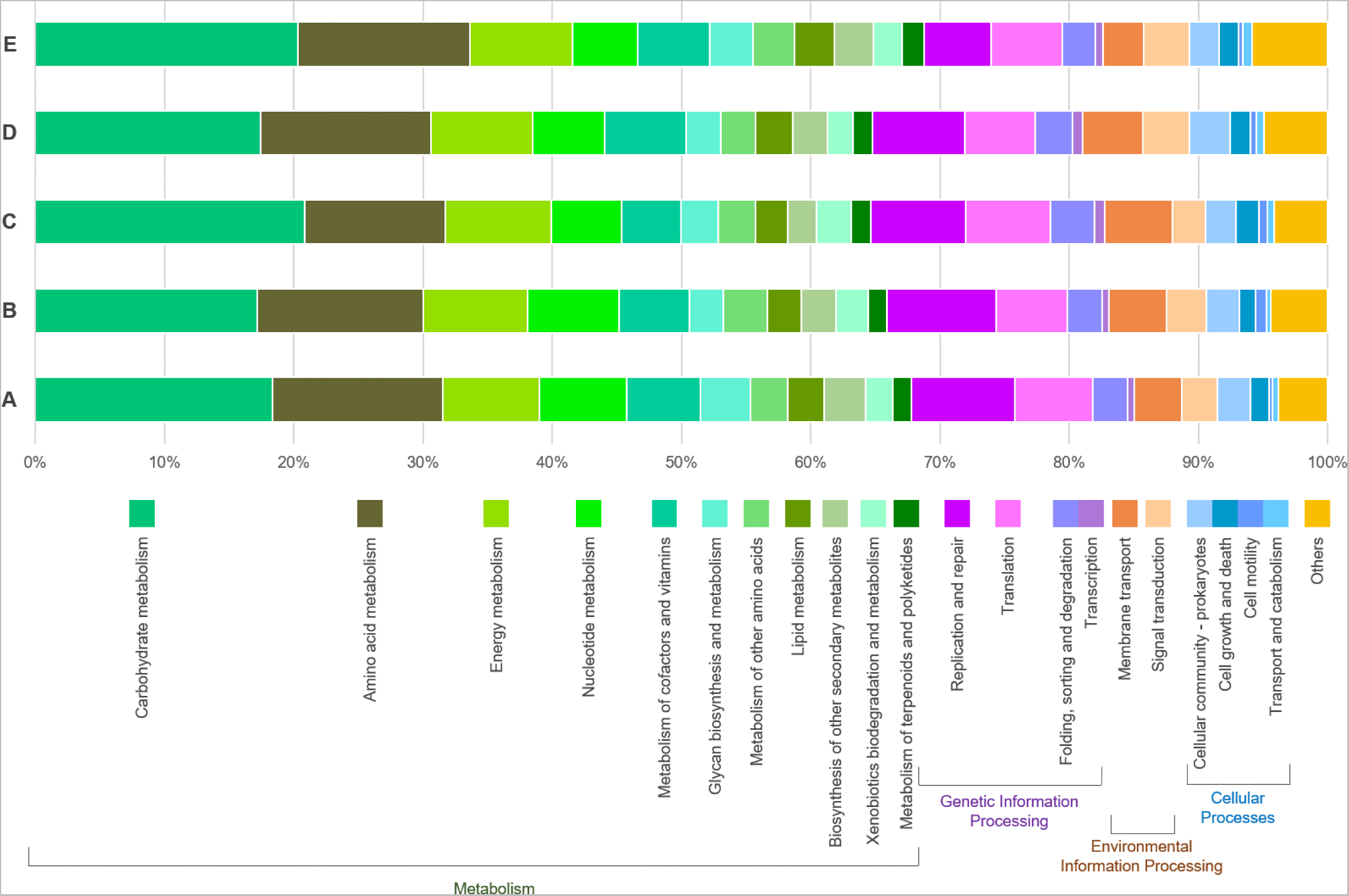
Functional characterization of the swine caecal microbiome according to the KEGG annotation pathway database at level 2.

Mining these genes at different KEGG Pathways help understand some of the specific metabolic processes that are important in the swine microbiome. Detailed descriptions of the putative function of identified genes at KEGG level 3 suggests that glycolysis/gluconeogenesis, citrate cycle (TCA cycle), pyruvate, starch and sucrose metabolism are key for bacterial survival in the swine microbiome; and were identified in samples from all farms. Similarly, genes involved in metabolic pathways for amino acid metabolism, including alanine, aspartate, glutamate, glycine, serine, threonine, cysteine, and methionine were also conserved across all samples and genes with roles in energy and nucleotide metabolism. In addition, the most common pathway among these farms was DNA replication, mismatch repair, and homologous recombination associated with replication and repair categories. When we considered genetic information processing, the aminoacyl-tRNA biosynthesis, and ribosome pathways were the most common across all farms. ABC transporters and two-component systems were also established as the most common pathway in the environmental information processing categories (**Figure 6**).

**Figure 6:**
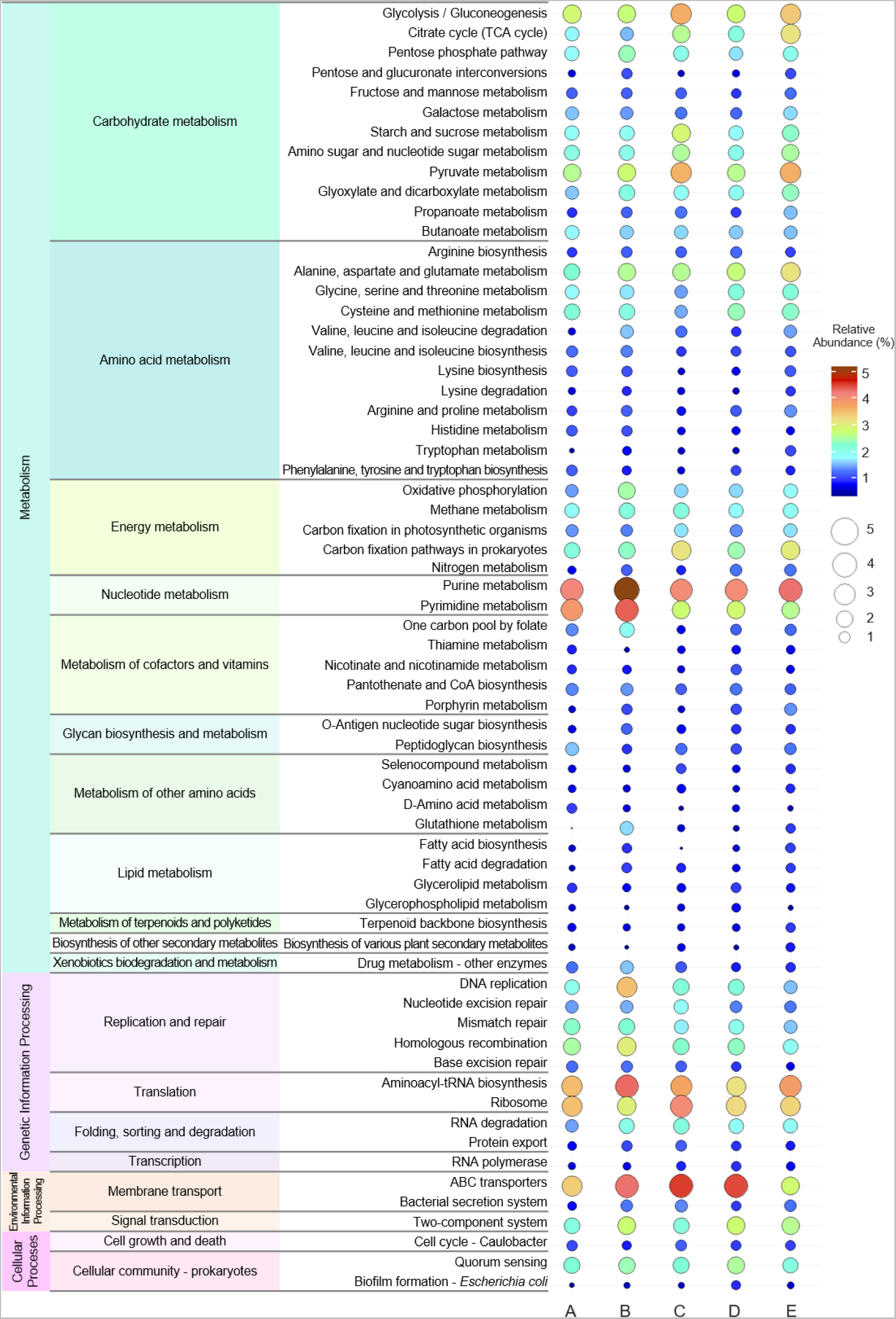
The balloon plot of the functional characterization of the swine caecal microbiome according to the KEGG annotation pathway database at level 3. Circle sizes and color represented the percentage of the relative abundance in each KEGG metabolic pathway.

### Antimicrobial resistome identification harboured in the swine commercial farms based on the caecal microbiome data

Antimicrobial Resistance Genes (ARGs) were detected through nucleotide comparisons with a curated list of ARGs in the CARD database (The Comprehensive Antibiotic Resistance Database). There were 30 ARGs detected among five caecal microbiomes, which putatively confer resistance against nine antimicrobial classes including aminoglycoside, beta-lactam, lincosamide, macrolide, macrolide-lincosamide-streptogramin B, nucleoside, sulfonamide, tetracycline, and trimethoprim (**Figure 7**). Genes that confer resistance to tetracycline were the most common ARG, with 15 genes identified from all farms, followed by aminoglycoside resistance (comprised 5 ARGS) and nucleoside resistance (comprised 3 ARGs). However, only one gene was found to be resistant to beta-lactam, lincosamide, macrolide, sulfonamide, and trimethoprim across all samples. The most common ARGs encountered in all farms were *APH(3’)-IIIa, tet(40),* and *tetW*, which leading to aminoglycoside and tetracycline resistance (**Figure 7**). For the number of ARGs detected in each farm, there are diverse ARGs in each farm, including farm E had the highest abundance of ARGs (8 ARGs), followed by farm A (7 ARGs), and farm D (6 ARGs). When focusing on each farm, some ARGs were detected in individual farms, including *InuC, ErmG* and *sul1* in farm A, which conferred to the lincosamide, macrolide-lindosamide-streptogramin B and sulfonamide resistance, respectively*. TetO*, which confers tetracycline resistance, and *dfrA12*, which encoded for trimethoprim resistance, were found in farms B and D, respectively. *CfxA2, mefA,* and *ErmF*, associated with beta-lactam, macrolide, and macrolide-lincosamide-streptogramin B resistance, were only found in farm E. Furthermore, we realized that some ARGs were shared among each farm, including *SAT-4* linked to the nucleoside resistance, which was shared among farm A, farm D, and farm E. *TetQ* was shared by farm C and farm E, whereas *tetA(P)* was shared by farm C and farm D. These two genes were encoded to tetracycline resistance (**Figure 7** & **Figure 8**).

**Figure 7:**
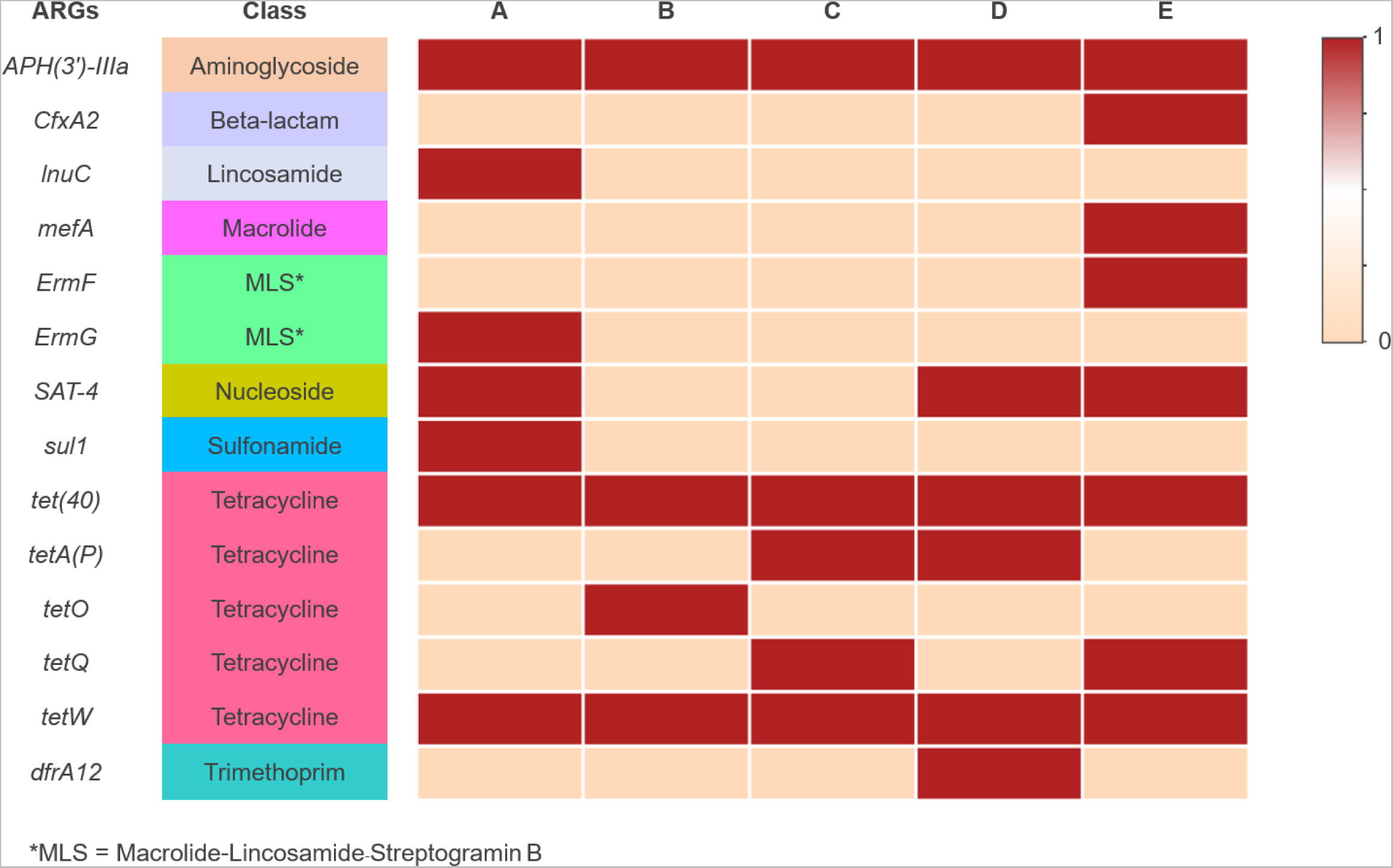
Binary heatmap analysis of antimicrobial resistance genes harbored in the swine caecal microbiome among five commercial farms.

**Figure 8:**
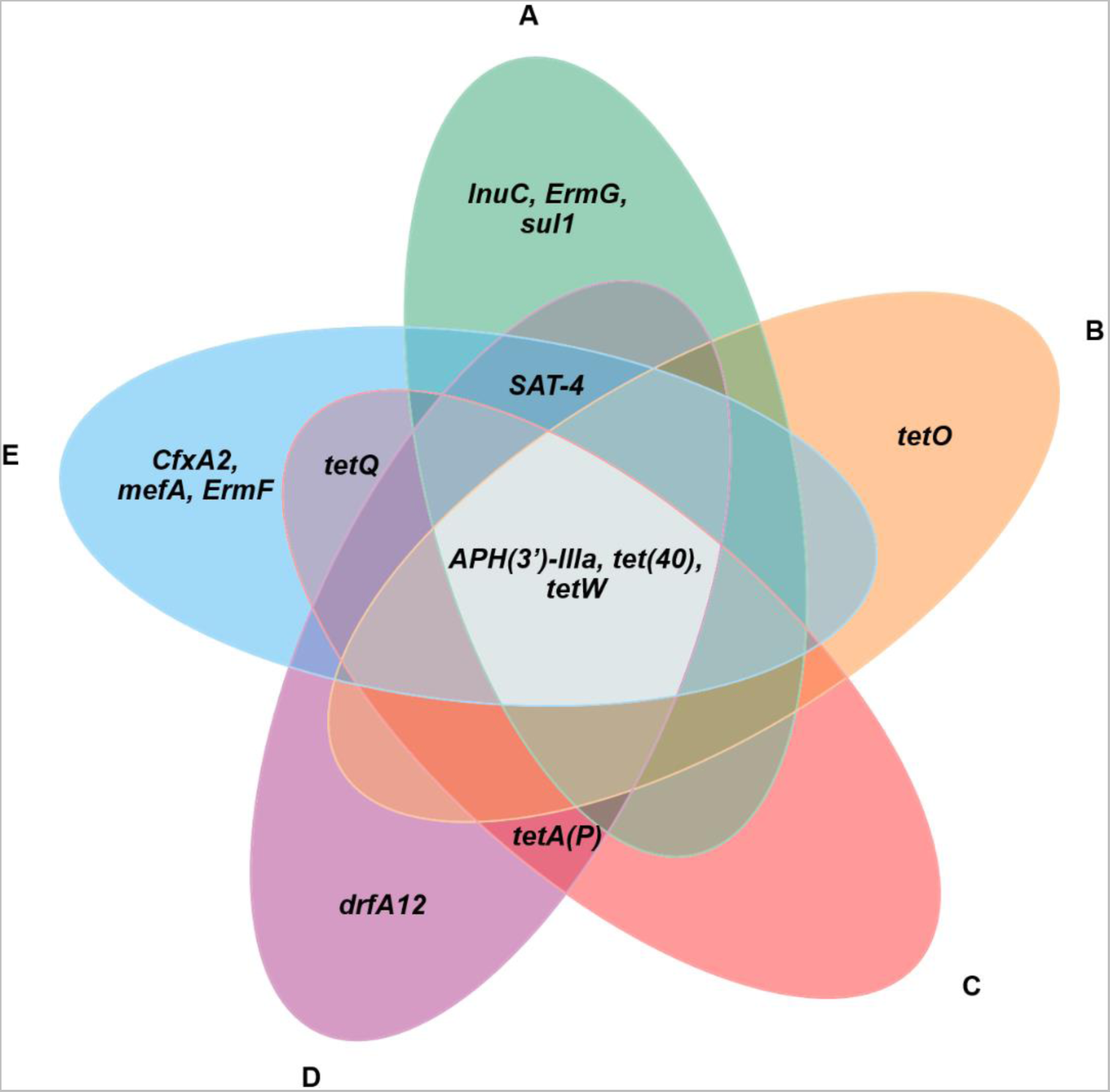
The Venn diagram of intersection analysis of antimicrobial resistance genes harbored in the swine caecal microbiome among five commercial farms.

## Discussion

Our study used shotgun metagenome sequencing to examine the diversity of microorganisms in the swine intestinal tract and their impact on physiology, health, and food safety by collecting samples using the intestinal scraping method. The bacterial composition varied in abundance and taxonomy across different swine farms due to various factors including farm location, size, swine type, and feed composition. Antimicrobial usage within the swine farm was associated with the heterogeneity of the bacterial composition found in the swine intestinal tract. Notably, the dominant bacteria differed among farms, including pathogenic, opportunistic, and commensal species, which correlate to different properties, both in terms of influencing the host’s physiology and causing disease in both humans and animals. Carriage of multiple species with the ability to cause disease demonstrated that swine could serve as a source of various pathogens beyond foodborne pathogens, which have adverse consequences for the livestock industry and public health (Looft et al., 2014; Guevarra et al., 2018; Chen et al., 2021).

The predominant phyla in the caecal mucosal epithelium samples were *Firmicutes*, *Actinobacteria*, *Proteobacteria*, and *Bacteroidota*, which is consistent with previous studies of swine caecal microbiomes (Yang et al., 2016; Tan et al., 2017; Quan et al., 2020). *Escherichia coli* was the predominant pathogen found in the caecal microbiome, which is a common pathogen in both humans and animals (McNally et al., 2016; Kim et al., 2022; Tiwari et al., 2023). This pathogen is associated with post-weaning diarrhea (PWD) and oedema disease in swine, which had an enormous economic impact. Moreover, *Escherichia coli* was the important cause of the foodborne illness in humans (Tseng et al., 2014). The high abundance of *Escherichia* genus correlated with antibiotic usage and affect to the bacteria homeostasis in the gastrointestinal tract of animals, leading to the development of antimicrobial resistance pathogens (Looft et al., 2014). *Clostridium botulinum* was identified in samples from every farm and commonly found in a variety of environments, such as, in soil, water, and the swine gastrointestinal tracts. This pathogen poses a specific food safety risk as its spore’s germination properties, which confer an ability to survive through processing stages and contaminate food product (Myllykoski et al., 2006). The *Corynebacterium* genus was the most common on farm A and is commonly found in animal manure and fertilizer used in agriculture. Interestingly, this pathogen was a commonly cause of human nosocomial infections (Salem et al., 2015). Furthermore, high prevalence of antimicrobial resistance are also observed in *Corynebacterium*, potentially correlated with the use of antibiotics in livestock (Xu et al., 2020; Song et al., 2021). Reports in Europe have helped characterize transmission events between farm workers and swine of specific livestock-associated lineages of *Staphylococcus aureus* (Anukool et al., 2011; Price et al., 2012; Ward et al., 2014). The symptoms range from a simple skin infection to the severity of a life-threatening disease among the elderly and immunocompromised people (Islam et al., 2020). These findings highlight the potential of swine to harbour disease-causing pathogens and the need for standardised sanitation practices on swine farms.

Commensal bacteria such as *Faecalibacterium*, *Lactobacillus*, *Bifidobacterium*, and *Streptomyces* - which are all beneficial for digestion, were identified in all swine caecal samples. *Faecalibacterium prausnitzii* is a commensal bacterium which is accountable for promoting healthy gut-associated butyrate production, enhancing feed efficiency and microbiome balance (Haenen et al., 2013). Furthermore, *Bifidobacterium* and *Lactobacillus* the lactic acid bacteria and qualify as probiotics, have the potential to improve feed utilisation, and control the balance of pathogenic bacteria in the gut (Yang et al., 2015). *Streptomyces* are common soil dwelling organisms with extensive ability to produce antimicrobial agents and useful secondary metabolites (Chater, 2016). Detection of this genus in our microbiome samples suggests a close relationship between the animals and the surrounding environment (Angelakis et al., 2013). *Collinsella* is found in both swine and human gastrointestinal tracts associated with plant-based diets and affects feed intake and efficiency in swine. Although its pathogenicity in swine is unclear, *Collinsella* is linked to important human diseases like irritable bowel syndrome (Malinen et al., 2010; Walker et al., 2011). Common bacterial species shared between swine and humans indicates a potential exchange of microbiomes, which may raise public health concerns (Islam et al., 2020; Duarte et al., 2021).

Metagenomic analysis of the swine caecal microbiome also gives us an opportunity to identify specific genes and functional processes that are important in the host-microbe relationship. Using comparisons with the KEGG database, we reconstructed metabolic pathways that were common among the microbiomes of our caecal samples. Functional pathways that were common among healthy swine microbiomes included those associated with processes of nutrient utilisation and metabolism, particularly those involved in carbohydrate and amino acid metabolism. Short-chain fatty acids produced through carbohydrate metabolism are essential for good intestinal health and feed efficiency (Chen et al., 2017; Quan et al., 2020). Amino acids like glutamate and aspartate are oxidative fuels of the intestine and are crucial for maintaining intestinal mucosal mass and integrity (Yang and Liao, 2019). ABC transporters and two-component system facilitate nutrient transport and the use of physiological substrates (Sonnenburg et al., 2006; Davidson et al., 2008; Quan et al., 2020). These findings highlight the significance of the caecal microbiome in swine health and the digestion process, however swine infected with pathogenic species also impairs digestion through changes effected by intestinal immune responses (Bin et al., 2018; Quan et al., 2020).

Food animals such as swine are regarded as one of the most important antimicrobial resistance pathogen reservoirs, and are exposed to extensive antibiotic usage, via widespread use of antimicrobials in livestock for disease prevention, production efficiency promotion, and treatment has led to the emergence of antimicrobial resistance. This global public health concern could result in over 10 million deaths from antimicrobial-resistant bacteria by 2050 if not addressed (Balouiri et al., 2016). Acquiring data from shotgun metagenomics is appropriate for an overview of the antimicrobial resistance genes within one environment which can provide evidence of long-term antimicrobial usage in the livestock industry (Gweon et al., 2019; Smith et al., 2023). The caecal microbiome from all five swine farms contained antimicrobial resistance genes for aminoglycoside and tetracycline resistance, which are commonly used in the swine industry (Lekagul et al., 2019). Various other groups of antimicrobial resistance genes, such as beta-lactam, lincosamide, macrolide, macrolide-lincosamide-streptogramin B, nucleoside, sulfonamide, and trimethoprim resistance genes, were also found across the farms. These findings indicate a strong selective pressure resulting from widespread antimicrobial use and highlight the abundance of antimicrobial resistance organisms in industrially farmed swine (Witte, 2000; Harada & Asai, 2010; Kittiwan et al., 2022). Rapid industrialisation of agricultural processes has diminished physical barriers to recombination, facilitating the spread of AMR genes between species and gene pools (Mourkas et al., 2022), often via the exchange of mobile genetic elements and horizontal gene transfer (Johnson et al., 2016; Vinayamohan et al., 2022). While efforts have been made to prohibit the use of certain antimicrobials for agricultural use, we still identified several ARGs in our swine caecal microbiome samples, consistent with other studies (Lekagul et al., 2021; Nuangmek et al., 2021). The use of swine manure in agriculture may also been encouraging the dissemination of ARGs from livestock to the environment (Lima et al., 2020).

Our study provides useful baseline details on the variation one should expect to see within and between swine microbiomes from different farm production environments. However, due to our limited sample size further studies will be required to investigate correlation between modifications to the production environment and swine gut health. The intestinal scraping method used in this study was prone to contamination, with a high relatively high number of reads attributed to the host species (*S. scrofa*). This resulted in fewer reads that could be used to interrogate the microbiome; however, the method did allow us to focus on bacteria specifically attached to the caecal epithelial cell (Kelly et al., 2017). Increased resolution of the microbiome will also facilitate assembly of a broader number of taxonomic groups and individual species. This can provide a window on organisms that have previously been unculturable (Schloss & Handelsman., 2005; Liu et al., 2022). Functional redundancy in metabolic pathways of organisms’ resident in this very specific niche impairs our ability to segregate and identify beneficial/pathogenic bacterial species (Zhu et al., 2022; Fachrul et al., 2022).

## Conclusion

Our investigation revealed a diverse array of bacteria inhabiting the swine’s caecal epithelium, including pathogens capable of causing diseases in both humans and animals. These findings underscore the potential risk of transmission of these pathogens from swine farms to humans, emphasising the need for standardised sanitation measures in swine production facilities. We identified commensal bacteria associated with swine feed utilisation and efficiency. By leveraging the KEGG metabolic pathway database, we established a link between these bacteria and healthy swine physiology and feed utilisation. This understanding can aid in the development of strategies to enhance swine health and optimise feed efficiency. Our study also identified antimicrobial resistance genes within the swine caecal microbiome. This finding raises concerns about the potential impact of historical antibiotic use in swine farms on the prevalence of antimicrobial resistance. Implementing measures to mitigate antimicrobial resistance is crucial for safeguarding public health and ensuring the effectiveness of antibiotics.

## Author Contributions

TE conceived and designed the experiments, performed the experiment, analysed the data, prepared figures and/or tables, and approved the final draft. PT conceived and designed the experiments, performed the experiment, authored or reviewed drafts of the paper, and approved the final draft. SB conceived and designed the experiments, analysed the data and authored or reviewed drafts of the paper. PC conceived and designed the experiments, performed the experiment, analysed the data, prepared figures and/or tables, and approved the final draft. NT conceived and designed the experiments, collected the samples, performed the experiment, authored or reviewed drafts of the paper. BP conceived and designed the experiments, analysed the data, authored or reviewed drafts of the paper, and approved the final draft. PP conceived and designed the experiments, analysed the data, prepared figures and/or tables, authored or reviewed drafts of the paper, and approved the final draft.

## Acknowledgments

The authors would like to gratefully thank our colleagues at Chiang Mai University for their contribution. Analysis was conducted on high performance computing clusters at the University of Oxford, University of Arizona and MRC CLIMB (funded by the Medical Research Council; MR/L015080/1 & MR/T030062/1).

---

**Supplementary Table S1:**
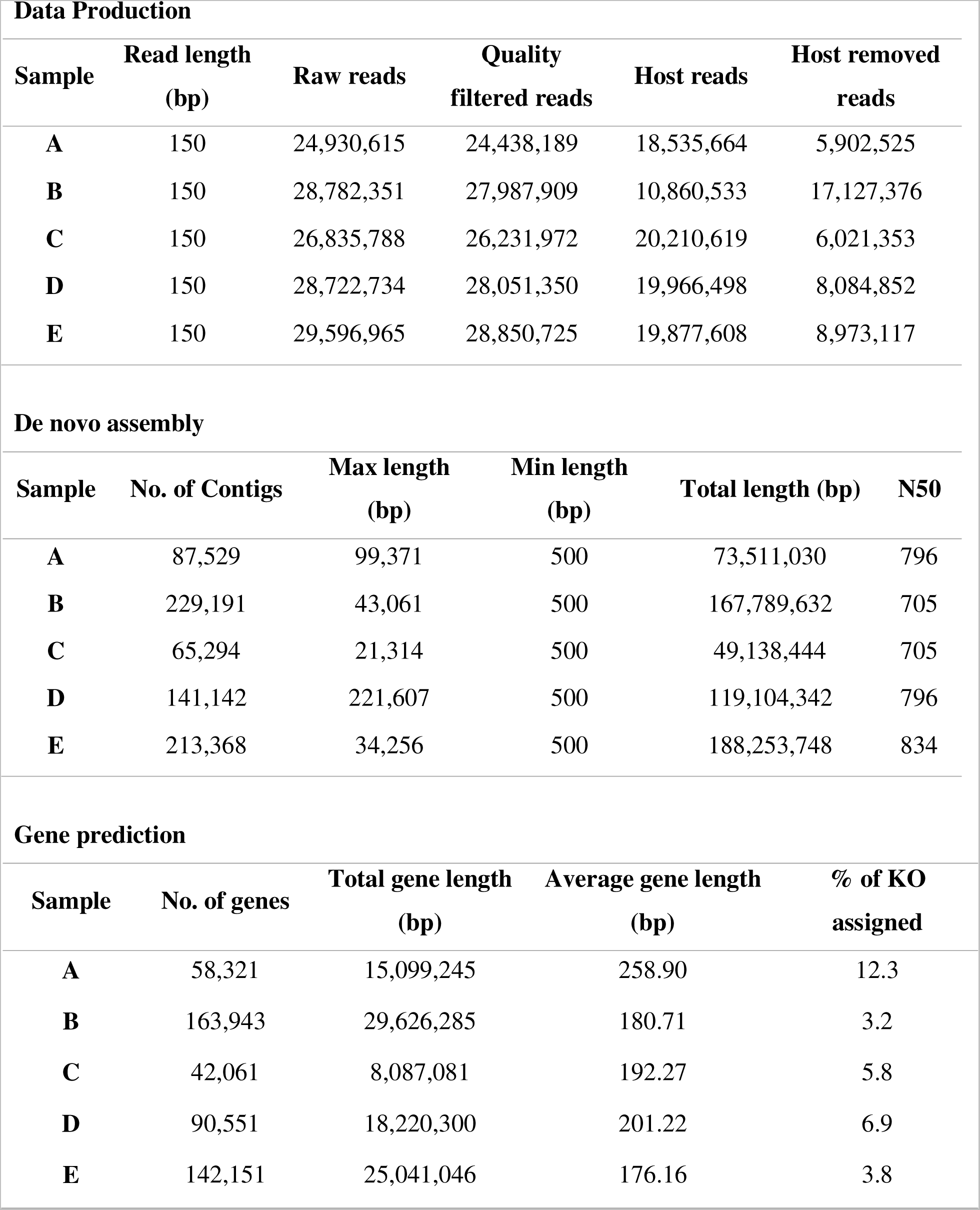
The summary of the metagenomic sequencing data and statistics for each sample.

**Supplementary Table S2:**
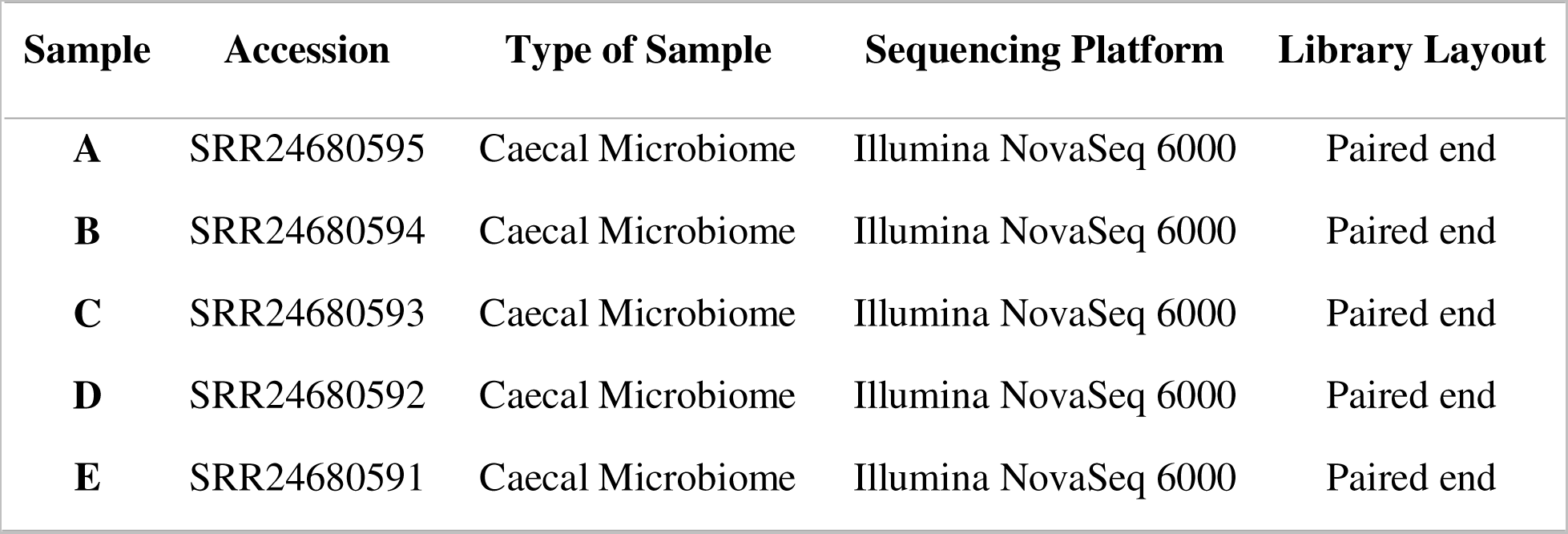
The individual accession number of the metagenomic sequencing data.

| **Sample** | **Accession** | **Type of Sample** | **Sequencing Platform** | **Library Layout** |
| --- | --- | --- | --- | --- |
| **A** | SRR24680595 | Caecal Microbiome | Illumina NovaSeq 6000 | Paired end |
| **B** | SRR24680594 | Caecal Microbiome | Illumina NovaSeq 6000 | Paired end |
| **C** | SRR24680593 | Caecal Microbiome | Illumina NovaSeq 6000 | Paired end |
| **D** | SRR24680592 | Caecal Microbiome | Illumina NovaSeq 6000 | Paired end |
| **E** | SRR24680591 | Caecal Microbiome | Illumina NovaSeq 6000 | Paired end |

Supplementary Figure S1: The relative abundance and diversity of the swine caecal microbiome at genus level distributed in each commercial farm.

Supplementary Figure S2: The relative abundance and diversity of the swine caecal microbiome at species level distributed in each commercial farm.

## References

Alcock, B.P., Raphenya, A.R., Lau, T.T., Tsang, K.K., Bouchard, M., Edalatmand, A., Huynh, W., Nguyen, A.L.V., Cheng, A.A., Liu, S. and Min, S.Y., 2020. CARD 2020: antibiotic resistome surveillance with the comprehensive antibiotic resistance database. Nucleic acids research, 48(D1), pp.D517–D525.

Andrews, S., Krueger, F., Segonds-Pichon, A., Biggins, L., Krueger, C. and Wingett, S., 2010. FastQC. A quality control tool for high throughput sequence data, 370.

Angelakis, E., Merhej, V. and Raoult, D., 2013. Related actions of probiotics and antibiotics on gut microbiota and weight modification. The Lancet infectious diseases, 13(10), pp.889–899.

Anukool, U., O’Neill, C.E., Butr-Indr, B., Hawkey, P.M., Gaze, W.H. and Wellington, E.M., 2011. Meticillin-resistant *Staphylococcus aureus* in pigs from Thailand. International journal of antimicrobial agents, 38(1), pp.86–87.

Apiwatsiri, P., Pupa, P., Sirichokchatchawan, W., Sawaswong, V., Nimsamer, P., Payungporn, S., Hampson, D.J. and Prapasarakul, N., 2022. Metagenomic analysis of the gut microbiota in piglets either challenged or not with enterotoxigenic Escherichia coli reveals beneficial effects of probiotics on microbiome composition, resistome, digestive function and oxidative stress responses. Plos one, 17(6), p.e0269959.

Balouiri, M., Sadiki, M. and Ibnsouda, S.K., 2016. Methods for in vitro evaluating antimicrobial activity: A review. Journal of pharmaceutical analysis, 6(2), pp.71–79.

Bin, P., Tang, Z., Liu, S., Chen, S., Xia, Y., Liu, J., Wu, H. and Zhu, G., 2018. Intestinal microbiota mediates Enterotoxigenic *Escherichia coli*-induced diarrhea in piglets. BMC veterinary research, 14(1), pp.1–13.

Chater, K.F., 2016. Recent advances in understanding *Streptomyces*. F1000Research, 5.

Chen, C., Zhou, Y., Fu, H., Xiong, X., Fang, S., Jiang, H., Wu, J., Yang, H., Gao, J. and Huang, L., 2021. Expanded catalog of microbial genes and metagenome-assembled genomes from the pig gut microbiome. Nature communications, 12(1), p.1106.

Chen, L., Xu, Y., Chen, X., Fang, C., Zhao, L. and Chen, F., 2017. The maturing development of gut microbiota in commercial piglets during the weaning transition. Frontiers in microbiology, 8, p.1688.

Danecek, P., Bonfield, J.K., Liddle, J., Marshall, J., Ohan, V., Pollard, M.O., Whitwham, A., Keane, T., McCarthy, S.A., Davies, R.M. and Li, H., 2021. Twelve years of SAMtools and BCFtools. Gigascience, 10(2), p.giab008.

Davidson, A.L., Dassa, E., Orelle, C. and Chen, J., 2008. Structure, function, and evolution of bacterial ATP-binding cassette systems. Microbiology and molecular biology reviews, 72(2), pp.317–364.

Department of Livestock and Development, 2023. Number of farmers and animal populations 2022. (Available online at: https://ict.dld.go.th/webnew/images/stories/stat_web/yearly/2565/province/T5-1-Pig.pdf, accessed February 6,2023).

Duarte, A.S.R., Röder, T., Van Gompel, L., Petersen, T.N., Hansen, R.B., Hansen, I.M., Bossers, A., Aarestrup, F.M., Wagenaar, J.A. and Hald, T., 2021. Metagenomics-based approach to source-attribution of antimicrobial resistance determinants–identification of reservoir resistome signatures. Frontiers in microbiology, 11, p.601407.

Fachrul, M., Méric, G., Inouye, M., Pamp, S.J. and Salim, A., 2022. Assessing and removing the effect of unwanted technical variations in microbiome data. Scientific Reports, 12(1), p.22236.

Fu, L., Niu, B., Zhu, Z., Wu, S. and Li, W., 2012. CD-HIT: accelerated for clustering the next-generation sequencing data. Bioinformatics, 28(23), pp.3150–3152.

Gweon, H.S., Shaw, L.P., Swann, J., De Maio, N., AbuOun, M., Niehus, R., Hubbard, A., Bowes, M.J., Bailey, M.J., Peto, T.E. and Hoosdally, S.J., 2019. The impact of sequencing depth on the inferred taxonomic composition and AMR gene content of metagenomic samples. Environmental Microbiome, 14(1), pp.1–15.

Guevarra, R.B., Hong, S.H., Cho, J.H., Kim, B.R., Shin, J., Lee, J.H., Kang, B.N., Kim, Y.H., Wattanaphansak, S., Isaacson, R.E. and Song, M., 2018. The dynamics of the piglet gut microbiome during the weaning transition in association with health and nutrition. Journal of animal science and biotechnology, 9, pp.1–9.

Haenen, D., Zhang, J., Souza da Silva, C., Bosch, G., van der Meer, I.M., van Arkel, J., van den Borne, J.J., Pérez Gutiérrez, O., Smidt, H., Kemp, B. and Müller, M., 2013. A diet high in resistant starch modulates microbiota composition, SCFA concentrations, and gene expression in pig intestine. The Journal of nutrition, 143(3), pp.274–283.

Harada, K. and Asai, T., 2010. Role of antimicrobial selective pressure and secondary factors on antimicrobial resistance prevalence in Escherichia coli from food-producing animals in Japan. Journal of biomedicine and biotechnology, 2010.

Hugenholtz, P., Goebel, B.M. and Pace, N.R., 1998. Impact of culture-independent studies on the emerging phylogenetic view of bacterial diversity. Journal of bacteriology, 180(18), pp.4765–4774.

Islam, M.Z., Johannesen, T.B., Lilje, B., Urth, T.R., Larsen, A.R., Angen, Ø. and Larsen, J., 2020. Investigation of the human nasal microbiome in persons with long-and short-term exposure to methicillin-resistant *Staphylococcus aureus* and other bacteria from the pig farm environment. Plos one, 15(4), p.e0232456.

Johnson, T.A., Stedtfeld, R.D., Wang, Q., Cole, J.R., Hashsham, S.A., Looft, T., Zhu, Y.G. and Tiedje, J.M., 2016. Clusters of antibiotic resistance genes enriched together stay together in swine agriculture. MBio, 7(2), pp.e02214–15.

Kanehisa, M., Sato, Y. and Morishima, K., 2016. BlastKOALA and GhostKOALA: KEGG tools for functional characterization of genome and metagenome sequences. Journal of molecular biology, 428(4), pp.726–731.

Kang, D.D., Li, F., Kirton, E., Thomas, A., Egan, R., An, H. and Wang, Z., 2019. MetaBAT 2: an adaptive binning algorithm for robust and efficient genome reconstruction from metagenome assemblies. PeerJ, 7, p.e7359.

Kelly, J., Daly, K., Moran, A.W., Ryan, S., Bravo, D. and ShirazilBeechey, S.P., 2017. Composition and diversity of mucosalassociated microbiota along the entire length of the pig gastrointestinal tract; dietary influences. Environmental microbiology, 19(4), pp.1425–1438.

Kim, H., Cho, J.H., Song, M., Cho, J.H., Kim, S., Kim, E.S., Keum, G.B., Kim, H.B. and Lee, J.H., 2021. Evaluating the prevalence of foodborne pathogens in livestock using metagenomics approach. Journal of Microbiology and Biotechnology, 31, 1701–8.

Kim, K., Song, M., Liu, Y. and Ji, P., 2022. Enterotoxigenic *Escherichia coli* infection of weaned pigs: Intestinal challenges and nutritional intervention to enhance disease resistance. Frontiers in Immunology, p.4370.

Kittiwan, N., Calland, J.K., Mourkas, E., Hitchings, M.D., Murray, S., Tadee, P., Tadee, P., Duangsonk, K., Meric, G., Sheppard, S.K. and Patchanee, P., 2022. Genetic diversity and variation in antimicrobial-resistance determinants of non-serotype 2 *Streptococcus suis* isolates from healthy pigs. Microbial Genomics, 8(11), p.000882.

Langmead, B. and Salzberg, S.L., 2012. Fast gapped-read alignment with Bowtie 2. Nature methods, 9(4), pp.357–359.

Lekagul, A., Tangcharoensathien, V. and Yeung, S., 2019. Patterns of antibiotic use in global pig production: a systematic review. Veterinary and animal science, 7, p.100058.

Lekagul, A., Tangcharoensathien, V., Liverani, M., Mills, A., Rushton, J. and Yeung, S., 2021. Understanding antibiotic use for pig farming in Thailand: a qualitative study. Antimicrobial Resistance & Infection Control, 10(1), pp.1–11.

Lima, T., Domingues, S. and Da Silva, G.J., 2020. Manure as a potential hotspot for antibiotic resistance dissemination by horizontal gene transfer events. Veterinary sciences, 7(3), p.110.

Liu, S., Moon, C.D., Zheng, N., Huws, S., Zhao, S. and Wang, J., 2022. Opportunities and challenges of using metagenomic data to bring uncultured microbes into cultivation. Microbiome, 10(1), p.76.

Looft, T., Allen, H.K., Cantarel, B.L., Levine, U.Y., Bayles, D.O., Alt, D.P., Henrissat, B. and Stanton, T.B., 2014. Bacteria, phages and pigs: the effects of in-feed antibiotics on the microbiome at different gut locations. The ISME journal, 8(8), pp.1566–1576.

Lu, J., Breitwieser, F.P., Thielen, P. and Salzberg, S.L., 2017. Bracken: estimating species abundance in metagenomics data. PeerJ Computer Science, 3, p.e104.

Malinen, E., Krogius-Kurikka, L., Lyra, A., Nikkilä, J., Jääskeläinen, A., Rinttilä, T., Vilpponen-Salmela, T., von Wright, A.J. and Palva, A., 2010. Association of symptoms with gastrointestinal microbiota in irritable bowel syndrome. World journal of gastroenterology: WJG, 16(36), p.4532.

Martin, M., 2011. Cutadapt removes adapter sequences from high-throughput sequencing reads. *EMBnet*. journal, 17(1), pp.10–12.

McNally, A., Oren, Y., Kelly, D., Pascoe, B., Dunn, S., Sreecharan, T., Vehkala, M., Välimäki, N., Prentice, M.B., Ashour, A. and Avram, O., 2016. Combined analysis of variation in core, accessory and regulatory genome regions provides a super-resolution view into the evolution of bacterial populations. PLoS genetics, 12(9), p.e1006280.

Mourkas, E., Yahara, K., Bayliss, S.C., Calland, J.K., Johansson, H., Mageiros, L., Muñoz-Ramirez, Z.Y., Futcher, G., Méric, G., Hitchings, M.D. and Sandoval-Motta, S., 2022. Host ecology regulates interspecies recombination in bacteria of the genus *Campylobacter*. Elife, 11, p.e73552.

Myllykoski, J., Nevas, M., Lindström, M. and Korkeala, H., 2006. The detection and prevalence of *Clostridium botulinum* in pig intestinal samples. International journal of food microbiology, 110(2), pp.172–177.

Nuangmek, A., Rojanasthien, S., Yamsakul, P., Tadee, P., Eiamsam-ang, T., Thamlikitkul, V., Tansakul, N., Suwan, M., Prasertsee, T., Chotinun, S. and Patchanee, P., 2021. Perspectives on antimicrobial use in pig and layer farms in thailand: legislation, policy, regulations and potential: https://doi.org/10.12982/VIS.2021.001. Veterinary Integrative Sciences, 19(1), pp.1–21.

Nurk, S., Meleshko, D., Korobeynikov, A. and Pevzner, P.A., 2017. metaSPAdes: a new versatile metagenomic assembler. Genome research, 27(5), pp.824–834.

Patchanee, P., Tanamai, P., Tadee, P., Hitchings, M.D., Calland, J.K., Sheppard, S.K., Meunsene, D., Pascoe, B. and Tadee, P., 2020. Whole-genome characterisation of multidrug resistant monophasic variants of *Salmonella Typhimurium* from pig production in Thailand. PeerJ, 8, p.e9700.

Price, L.B., Stegger, M., Hasman, H., Aziz, M., Larsen, J., Andersen, P.S., Pearson, T., Waters, A.E., Foster, J.T., Schupp, J. and Gillece, J., 2012. *Staphylococcus aureus CC398*: host adaptation and emergence of methicillin resistance in livestock. MBio, 3(1), pp.e00305–11.

Quan, J., Wu, Z., Ye, Y., Peng, L., Wu, J., Ruan, D., Qiu, Y., Ding, R., Wang, X., Zheng, E. and Cai, G., 2020. Metagenomic characterization of intestinal regions in pigs with contrasting feed efficiency. Frontiers in microbiology, 11, p.32.

Salem, N., Salem, L., Saber, S., Ismail, G. and Bluth, M.H., 2015. *Corynebacterium urealyticum*: a comprehensive review of an understated organism. Infection and drug resistance, pp.129–145.

Schloss, P.D. and Handelsman, J., 2005. Metagenomics for studying unculturable microorganisms: cutting the Gordian knot. Genome biology, 6(8), pp.1–4.

Seemann, T., 2016. ABRicate: mass screening of contigs for antibiotic resistance genes. San Francisco, CA: .https://github.com/tseemann/abricate

Smith, R.P., May, H.E., AbuOun, M., Stubberfield, E., Gilson, D., Chau, K.K., Crook, D.W., Shaw, L.P., Read, D.S., Stoesser, N. and Vilar, M.J., 2023. A longitudinal study reveals persistence of antimicrobial resistance on livestock farms is not due to antimicrobial usage alone. Frontiers in Microbiology, 14, p.700.

Song, T., Li, H., Li, B., Yang, J., Sardar, M.F., Yan, M., Li, L., Tian, Y., Xue, S. and Zhu, C., 2021. Distribution of antibiotic-resistant bacteria in aerobic composting of swine manure with different antibiotics. Environmental Sciences Europe, 33(1), pp.1–13.

Sonnenburg, E.D., Sonnenburg, J.L., Manchester, J.K., Hansen, E.E., Chiang, H.C. and Gordon, J.I., 2006. A hybrid two-component system protein of a prominent human gut symbiont couples glycan sensing in vivo to carbohydrate metabolism. Proceedings of the National Academy of Sciences, 103(23), pp.8834–8839.

Strachan, C.R., Yu, X.A., Neubauer, V., Mueller, A.J., Wagner, M., Zebeli, Q., Selberherr, E. and Polz, M.F., 2023. Differential carbon utilization enables co-existence of recently speciated Campylobacteraceae in the cow rumen epithelial microbiome. Nature Microbiology, pp.1–12.

Tan, Z., Yang, T., Wang, Y., Xing, K., Zhang, F., Zhao, X., Ao, H., Chen, S., Liu, J. and Wang, C., 2017. Metagenomic analysis of cecal microbiome identified microbiota and functional capacities associated with feed efficiency in landrace finishing pigs. Frontiers in microbiology, 8, p.1546.

Team, R., RStudio: Integrated Development Environment for R (RStudio, PBC, Boston, MA, 2020). URL http://www.rstudio.com.

The Swine Raisers Association of Thailand, 2023. Livestock pig production and market trend 2023. (Available online at: https://www.swinethailand.com/17369771/livestock-pig-production-and-market-trend-2023, accessed February 6,2023).

Tiwari, S.K., van der Putten, B.C., Fuchs, T.M., Vinh, T.N., Bootsma, M., Oldenkamp, R., La Ragione, R., Matamoros, S., Hoa, N.T., Berens, C. and Leng, J., 2023. Genome-wide association reveals host-specific genomic traits in *Escherichia coli*. BMC biology, 21(1), p.76.

Tseng, M., Fratamico, P.M., Manning, S.D. and Funk, J.A., 2014. Shiga toxin-producing *Escherichia coli* in swine: the public health perspective. Animal health research reviews, 15(1), pp.63–75.

Tunsagool, P., Mhuantong, W., Tangphatsornruang, S., Am-In, N., Chuanchuen, R., Luangtongkum, T. and Suriyaphol, G., 2021. Metagenomics of antimicrobial and heavy metal resistance in the cecal microbiome of fattening pigs raised without antibiotics. Applied and Environmental Microbiology, 87(8), pp.e02684–20.

Vinayamohan, P.G., Pellissery, A.J. and Venkitanarayanan, K., 2022. Role of horizontal gene transfer in the dissemination of antimicrobial resistance in food animal production. Current Opinion in Food Science, 47, p.100882.

Walker, A.W., Ince, J., Duncan, S.H., Webster, L.M., Holtrop, G., Ze, X., Brown, D., Stares, M.D., Scott, P., Bergerat, A. and Louis, P., 2011. Dominant and diet-responsive groups of bacteria within the human colonic microbiota. The ISME journal, 5(2), pp.220–230.

Ward, M.J., Gibbons, C.L., McAdam, P.R., Van Bunnik, B.A.D., Girvan, E.K., Edwards, G.F., Fitzgerald, J.R. and Woolhouse, M.E.J., 2014. Time-scaled evolutionary analysis of the transmission and antibiotic resistance dynamics of *Staphylococcus aureus* clonal complex 398. Applied and environmental microbiology, 80(23), pp.7275–7282.

Witte, W., 2000. Selective pressure by antibiotic use in livestock. International journal of antimicrobial agents, 16, pp.19–24.

Wood, D.E., Lu, J. and Langmead, B., 2019. Improved metagenomic analysis with Kraken 2. Genome biology, 20, pp.1–13.

World Organization for Animal Health, 2023. One Health. (Available online at: https://www.woah.org/en/what-we-do/global-initiatives/one-health/, accessed February 6,2023).

Xiao, L., Estellé, J., Kiilerich, P., Ramayo-Caldas, Y., Xia, Z., Feng, Q., Liang, S., Pedersen, A.Ø., Kjeldsen, N.J., Liu, C. and Maguin, E., 2016. A reference gene catalogue of the pig gut microbiome. Nature microbiology, 1(12), pp.1–6.

Xu, Y., Li, H., Shi, R., Lv, J., Li, B., Yang, F., Zheng, X. and Xu, J., 2020. Antibiotic resistance genes in different animal manures and their derived organic fertilizer. Environmental Sciences Europe, 32, pp.1–10.

Yang, F., Hou, C., Zeng, X. and Qiao, S., 2015. The use of lactic acid bacteria as a probiotic in swine diets. Pathogens, 4(1), pp.34–45.

Yang, H., Huang, X., Fang, S., Xin, W., Huang, L. and Chen, C., 2016. Uncovering the composition of microbial community structure and metagenomics among three gut locations in pigs with distinct fatness. Scientific reports, 6(1), p.27427.

Yang, Z. and Liao, S.F., 2019. Physiological effects of dietary amino acids on gut health and functions of swine. Frontiers in veterinary science, 6, p.169.

Zhu, W., Lomsadze, A. and Borodovsky, M., 2010. Ab initio gene identification in metagenomic sequences. Nucleic acids research, 38(12), pp. e132–e132.

Zhu, Q., Huang, S., Gonzalez, A., McGrath, I., McDonald, D., Haiminen, N., Armstrong, G., Vázquez-Baeza, Y., Yu, J., Kuczynski, J. and Sepich-Poore, G.D., 2022. Phylogeny-aware analysis of metagenome community ecology based on matched reference genomes while bypassing taxonomy. Msystems, 7(2), pp.e00167–22.

Zou, A., Nadeau, K., Xiong, X., Wang, P.W., Copeland, J.K., Lee, J.Y., Pierre, J.S., Ty, M., Taj, B., Brumell, J.H. and Guttman, D.S., 2022. Systematic profiling of the chicken gut microbiome reveals dietary supplementation with antibiotics alters expression of multiple microbial pathways with minimal impact on community structure. Microbiome, 10(1), pp.1–29.

