## Supplementary Figure S1 for "Investigation of swine caecal microbiomes in the northern region of Thailand"

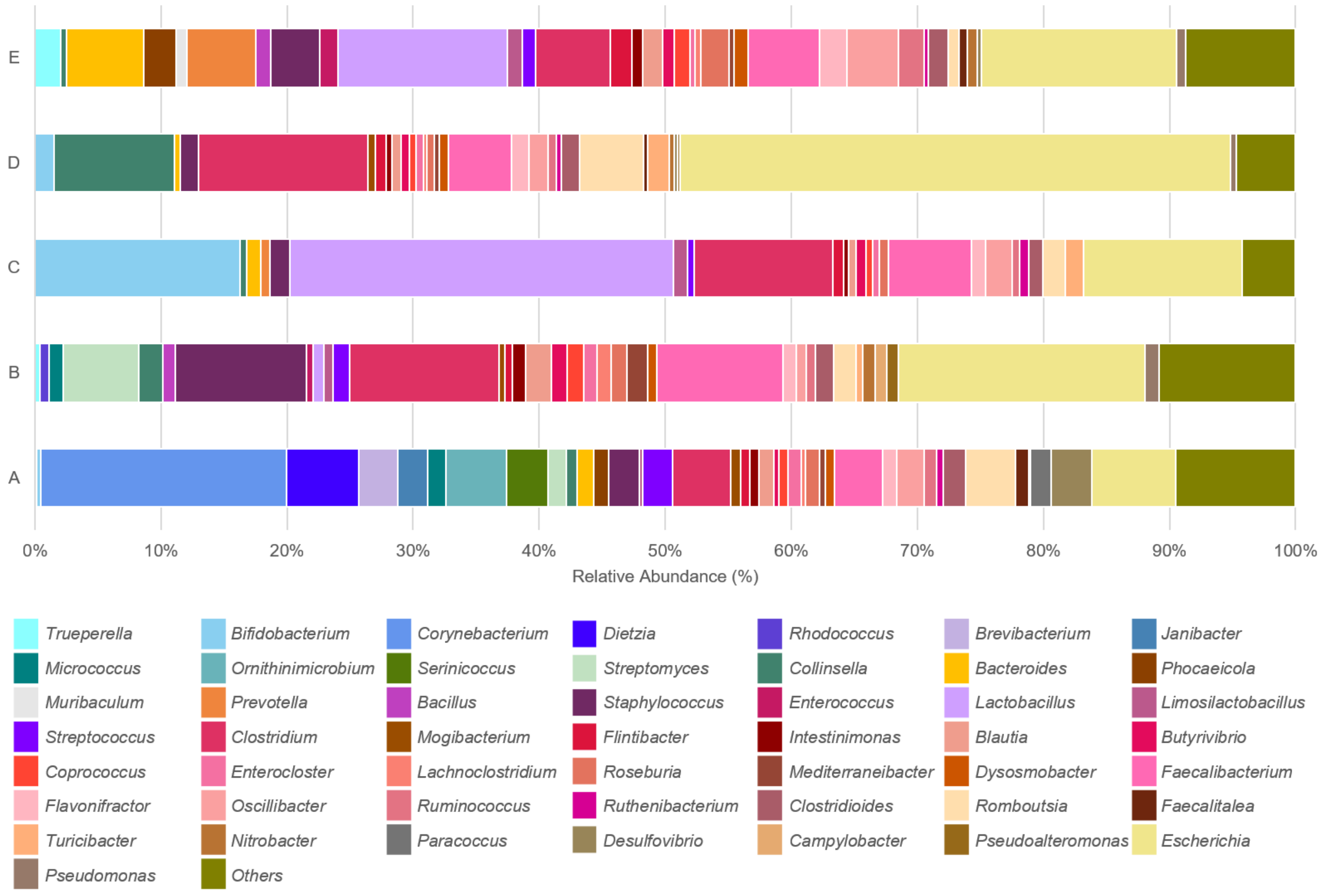


Supplementary Figure S1: The relative abundance and diversity of the swine caecal microbiome at genus level distributed in each commercial farm.
