## Supplementary Figure S2 for "Investigation of swine caecal microbiomes in the northern region of Thailand"

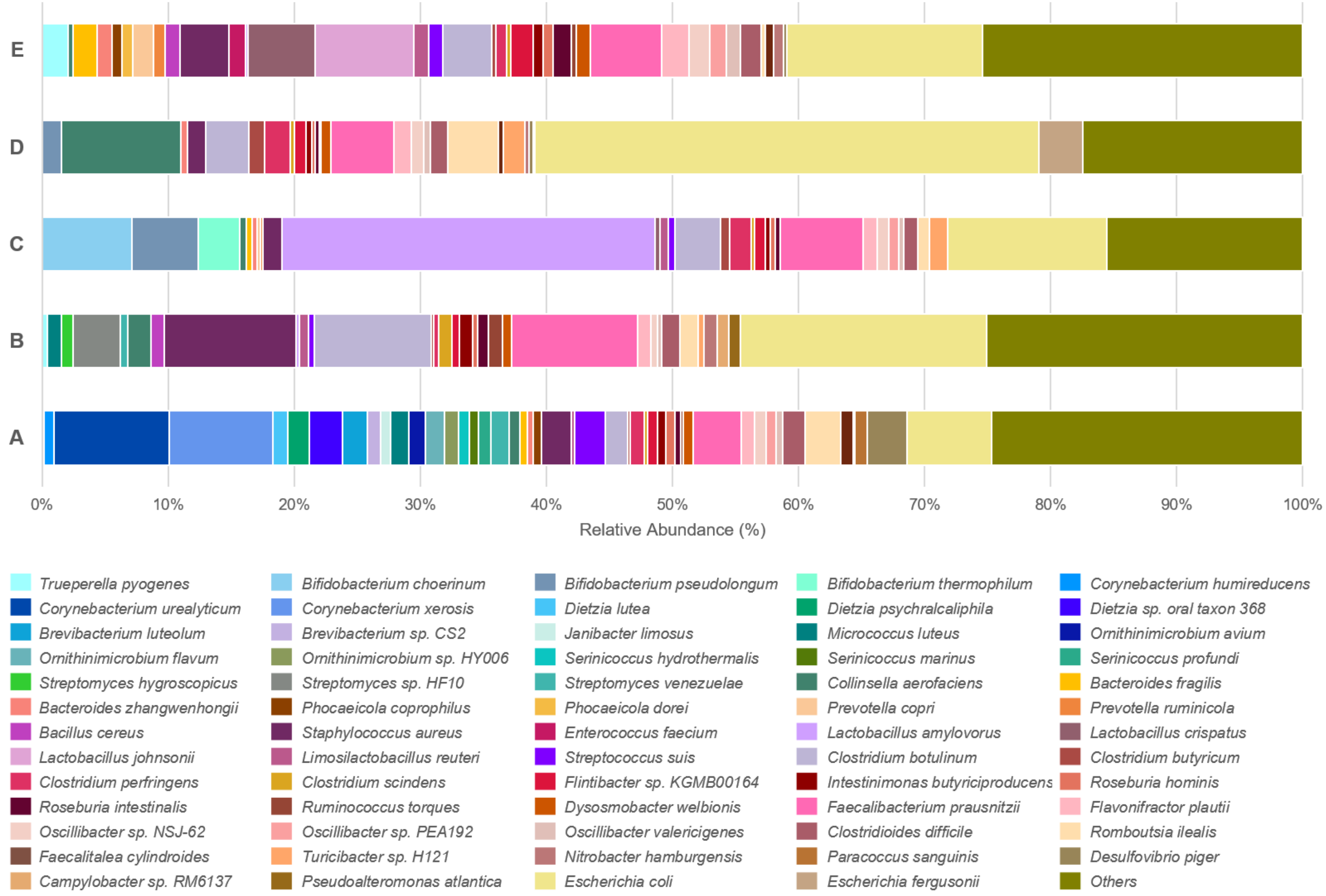


Supplementary Figure S2: The relative abundance and diversity of the swine caecal microbiome at species level distributed in each commercial farm.
